# The power of single molecule real-time sequencing technology in the de *novo* assembly of a eukaryotic genome

**DOI:** 10.1101/021634

**Authors:** Hiroaki Sakai, Ken Naito, Eri Ogiso-Tanaka, Yu Takahashi, Kohtaro Iseki, Chiaki Muto, Kazuhito Satou, Kuniko Teruya, Akino Shiroma, Makiko Shimoji, Takashi Hirano, Takeshi Itoh, Akito Kaga, Norihiko Tomooka

## Abstract

Second-generation sequencers (SGS) have been game-changing, achieving cost-effective whole genome sequencing in many non-model organisms. However, a large portion of the genomes still remains unassembled. We reconstructed azuki bean (*Vigna angularis*) genome using single molecule real-time (SMRT) sequencing technology and achieved the best contiguity and coverage among currently assembled legume crops. The SMRT-based assembly produced 100 times longer contigs with 100 times smaller amount of gaps compared to the SGS-based assemblies. A detailed comparison between the assemblies revealed that the SMRT-based assembly enabled a more comprehensive gene annotation than the SGS-based assemblies where thousands of genes were missing or fragmented. A chromosome-scale assembly was generated based on the high-density genetic map, covering 86% of the azuki bean genome. We demonstrated that SMRT technology, though still needed support of SGS data, achieved a near-complete assembly of a eukaryotic genome.

---

Genome projects used to consume a large amount of funds and labor. For example, the rice genome project^1^ took 14 years and cost several hundred million dollars. The paradigm was changed by the advent of pyrosequencing^2^ and Solexa sequencing^3^ technologies. The high-throughput sequencing capacity of these second-generation sequencers (SGS) enabled the assembly of diploid plant genomes with much less time and cost^4^.

However, the read length of SGS is not long enough to span repetitive sequences, which often comprise 50–80% of non-model plant genomes^5^. Although paired reads with long inserts could help resolve such repeated sequences, missing and fragmentation of gene coding sequences have been claimed^6,7^. As such, evolutionary studies based on such incorrect assemblies could reach incorrect conclusions. Moreover, simple misassemblies or mis-scaffolding could be deleterious in map-based cloning. Therefore, read length is one of the most important factors in determining the complete genome sequences.

The third generation, single molecule real-time (SMRT) sequencing platform^8^ now successfully generates reads of 10 kb on average^9^ and recently achieved an N50 of 4.3 Mb in assembling the haploid human genome^10^.

In *de novo* assembly, where a reference genome is not available, having a high-density genetic linkage map is also important. To reconstruct pseudomolecules of chromosomes, the assembled contigs/scaffolds have to be assigned according to the order of the marker loci. However, if the markers are not dense enough or evenly distributed, a large portion of the assembly can remain unanchored. In many cases, only 30–60% of the genomes have been assigned to pseudomolecules^11-19^.

Here, we present a near-complete genome sequence of the azuki bean (*Vigna angularis*), the second-most important grain legume in East Asia^20^. Nowadays, the breeding of azuki bean is extensively conducted and is targeting seed quality, cold tolerance, and disease resistance. However, the narrow genetic diversity of this domesticated species and the lack of high-quality genome sequences have limited the process. Although this species was recently sequenced, the draft assembly covered ~70% of the genome, and only half of it was anchored onto pseudomolecules^14^. As such, we sequenced the azuki bean genome using SMRT sequencing technology, in addition to SGS.

We tested several assembly approaches and found SMRT sequencing provided, by far, the best assembly. We also developed a high-density genetic map with evenly distributed markers, which we used not only for anchoring, but also for evaluating the accuracy of the assemblies. In addition, we evaluated the genome assemblies of legume crops based on some criteria used in Assemblathon 2^21^.

## RESULTS

### Genome sequencing using SGS

We obtained sequence data from the azuki bean, cultivar “Shumari”, using Roche and Illumina platforms. The details of sequencing libraries are shown in Supplementary Table 1. The k-mer distribution (k = 25) of our data indicated that the genome size of this cultivar was 540 Mb, which was a little larger than the C-value-based estimation (0.55/C = 531 Mb)^22^.

We first carried out a hybrid *de novo* assembly using both Roche and Illumina data to achieve the highest coverage as possible (Supplementary Fig. 1). We obtained 42,291 contigs, covering 84.0% of the genome, with an N50 size of kb. We then performed scaffolding and gap-closing and obtained 8,910 scaffolds, covering 93.5% of the genome, with an N50 size of 612.4 kb (see Assembly_1 in Table 1).

**Table 1.**
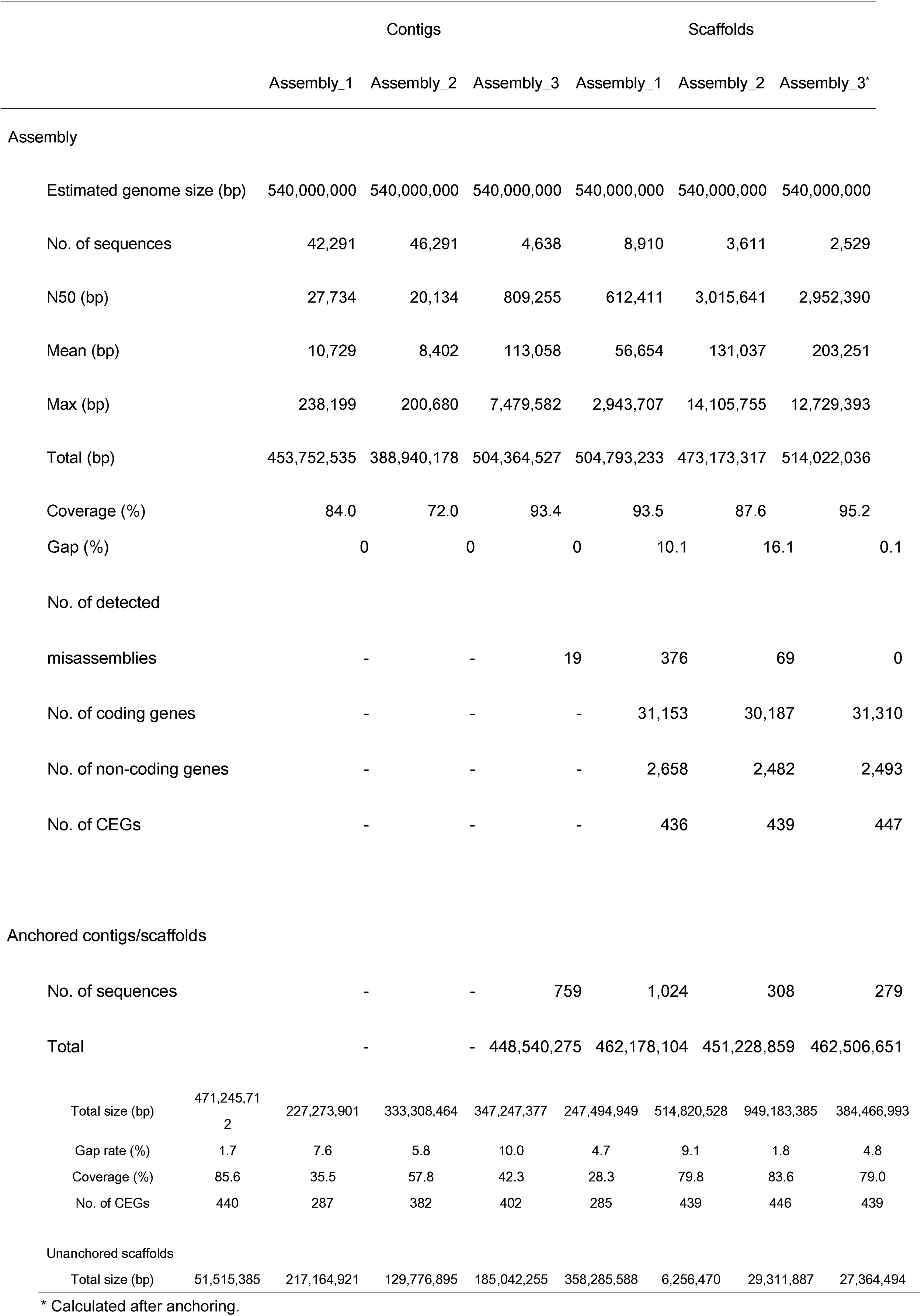
Statistics of the azuki genome assemblies.

We also tested an Illumina-only approach using ALLPATHS-LG^23^ (Supplementary Fig. 1). We obtained 46,291 contigs in 3,611 scaffolds, covering 72.0% and 87.6% of the genome, respectively. Although the total length of this assembly was smaller than that of Assembly_1, the N50 size and maximum length of the scaffolds were about five times larger (see Assembly_2 in Table 1).

### Construction of a high-density genetic map

To anchor the scaffolds onto the pseudomolecules, we developed a high-density genetic linkage map. We resequenced the genome of *V. nepalensis* which is a wild relative of the azuki bean (Supplementary Table 1), and developed a SNP array where the designed SNP markers were at least 100 kb away from each other. At the same time, we crossed *V. nepalensis* to the azuki bean and obtained 995 F2 plants. Genotyping the F2 plants with the SNP array resulted in a genetic map with 4,912 markers integrated into 11 linkage groups (LGs). The average marker distance was 0.12 cM, and the largest gap was 3.0 cM on LG9 (Supplementary Fig. 2).

### Anchoring and validation of SGS assemblies

According to the order of SNP markers in the linkage map, we assigned the scaffolds of the two assemblies. Of the 8,910 scaffolds of Assembly_1, 1,024 were anchored, covering 462 Mb (85.6%) of the azuki bean genome. However, we found 376 contradictions between the marker orders in the assembly and in the linkage map (Table 1). It indicated more than one-third of the anchored scaffolds were expected to contain assembly errors.

As for Assembly_2, only 308 scaffolds were anchored, but they covered 451 Mb (83.6% of the genome). Moreover, the anchored scaffolds contained much fewer contradictions than Assembly_1. However, more than one out of five among the anchored scaffolds still contained misassemblies.

### Assembly of long reads

Assembly errors could be deleterious for application of the genome sequence, including marker-assisted selection or map-based gene cloning. As such, we decided to adopt SMRT sequencing technology. The data we obtained had about 51× coverage of the azuki bean genome, with the average and the longest read length of 5.4 kb and 39.4 kb, respectively (Supplementary Table 1). Assembling the PacBio reads (Supplementary Fig. 3) resulted in 4,638 contigs covering 504 Mb or 93.4% of the azuki bean genome (see Assembly_3 in Table 1). The N50 and the maximum length of the contigs were 809 kb and 7.5 Mb, respectively, both of which were almost 30 times larger than those of SGS contigs.

### Validation of the long read assembly

We could anchor 759 of the 4,638 contigs onto the linkage map, covering 448 Mb (83.1% of the genome) with only 19 contradictions (Supplementary Fig. 3, Table 1). All misassembly sites were manually confirmed by mapping Illumina reads onto the contigs (Supplementary Fig. 4).

By mapping short reads, we also found conflicts between Assembly_3 and the short reads at the nucleotide level. Of these, 1,631 were substitutions, and 8,611 and 38,889 were insertions and deletions, respectively (Supplementary Fig. 5). The indels were greatly dominated by homopolymer sites of more than three consecutive nucleotides. We randomly chose 91 indel sites to validate by Sanger sequencing, and found that all indels in the short reads were correct (Supplementary Table 2), although all substitutions sites were in repetitive sequences and thus were impossible to validate. As such, we corrected all the detected indels according to the short reads (Supplementary Fig. 3).

To further validate the nucleotide-level accuracy of PacBio assembly, we obtained data from PacBio data release (https://github.com/PacificBiosciences/DevNet/wiki/) on *Arabidopsis thaliana* (L*er*-0) and *Drosophila melanogaster*. In both cases, errors of insertions/deletions at homopolymer sites dominated any other types of errors (Supplementary Fig. 5).

### Final assembly process on Assembly_3

After all these steps described above, we performed final scaffolding with Illumina’s paired-end reads with long inserts, excluding contradictions against the linkage map. As a result, we found 297 gaps in the scaffolds, where the overhangs of the bridged contigs were longer than the expected insert size. However, aligning two such bridged contigs to each other revealed, in 226 cases, that the terminal sequences of the two contigs overlapped and thus were merged. In addition, we performed gap-filling and closed 705 gaps.

The final assembly was comprised of 2,529 scaffolds covering 514 Mb (95.2%) of the genome, with an N50 of 3.0 Mb and a gap fraction of only 0.12%. Of these, 279 scaffolds were anchored, covering 462 Mb (85.6% of the genome) (Table 1). To reconstruct the pseudomolecules, we filled the gaps between scaffolds with the estimated number of Ns between the neighboring scaffolds. The resulting pseudomolecules ranged from 28.9 Mb (LG10) to 67.1 Mb (LG1) in size (Supplementary Table 3). All the pseudomolecules had telomeric repeat-like sequences (TTTAGGG) in both termini, except LG5, 6, and 11, where a telomeric repeat was found only in one end. The gap amount of the pseudomolecules was 1.7% (Table 2).

**Table 2.**
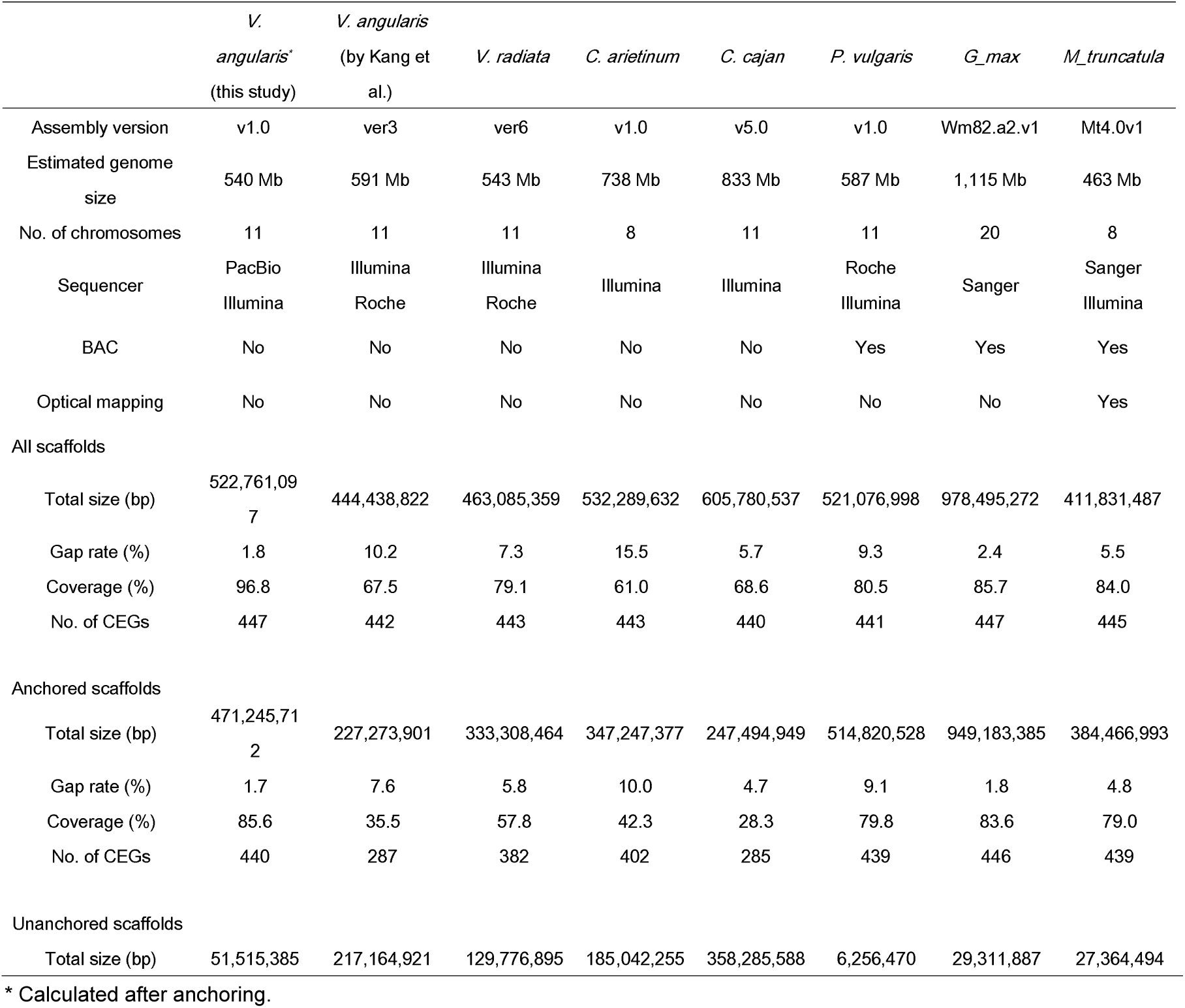
Statistics of recently assembled legume genomes.

In addition to N50 values, we calculated NG values (NG1 through NG100) where NG50, for example, is determined by taking the last-counted contig/scaffold size over the sum of all contig/scaffold sizes, from the longest to the shortest, until the sum reaches 50% of the estimated genome size^21^. The resulting “NG graph” can visualize differences in contig/scaffold lengths and coverage between the assemblies^21^ (Fig. 1).

As shown in Figure 1a, the scaffold NG graph of Assembly_2 was almost the same as that of Assembly_3 until NG80, whereas that of Assembly_1 was much lower. However, in the contig NG graph, Assembly_2 was 60–80 times lower than Assembly_3 and more than 100 times lower around NG 70 (Fig. 1b). Assembly_1 was better than Assembly_2 only in the context of coverage (Fig. 1b).

**Figure 1.**
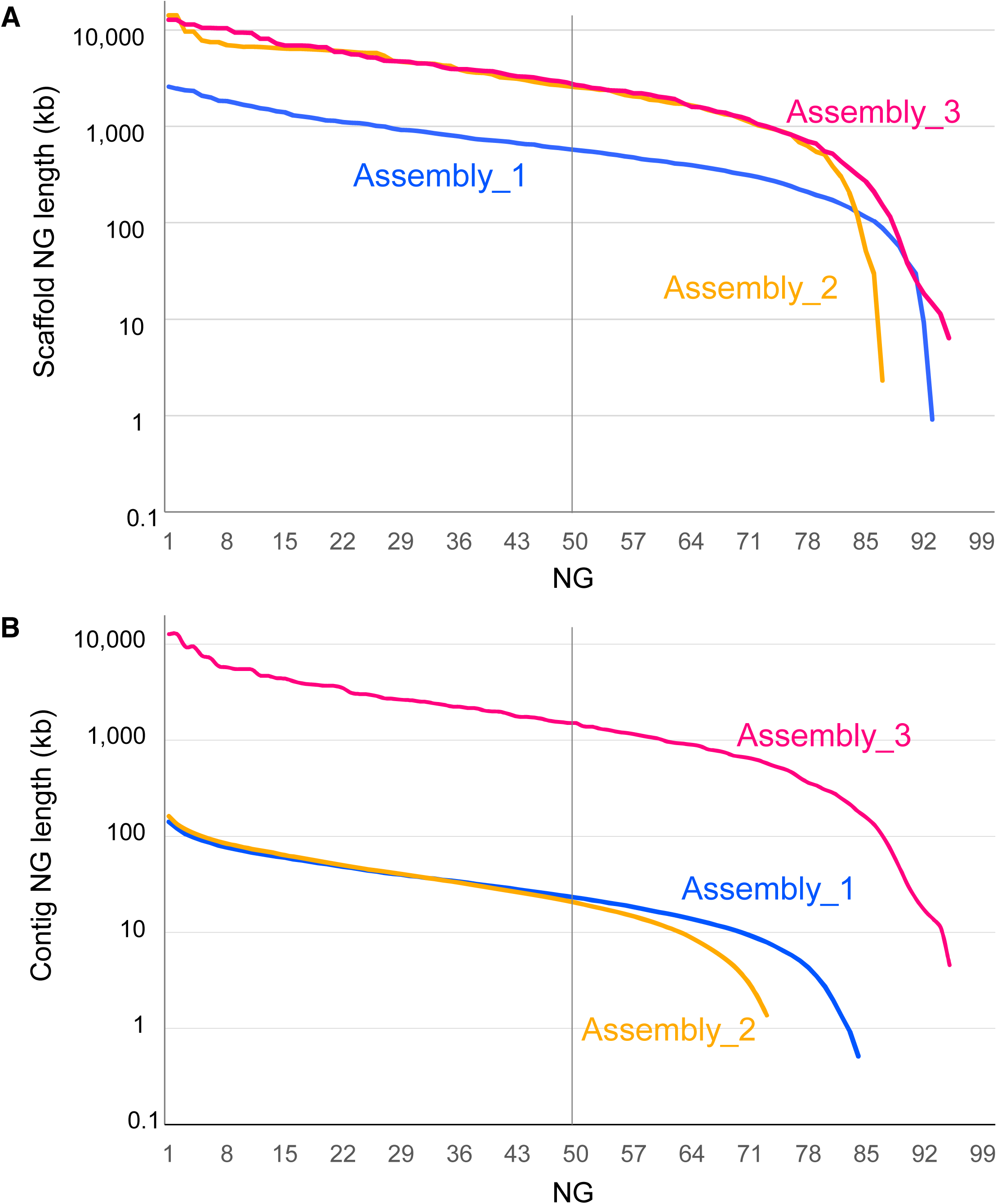
NG graphs of the three assemblies in scaffold length (a) and contig length (b). The y-axis indicates the calculated NG contig length (NG1 through NG100, see text for detail) in each assembled genome. The vertical line indicates the NG50 contig length.

### Annotation

Before gene annotation, we identified repeat elements to mask the assembled sequences. As expected, the amounts of repeats were the largest in Assembly_3 and the smallest in Assembly_2 (Fig. 2a). Of the estimated genome size, Assembly_3 had 273 Mb (50.6%) as repeat-masked, whereas Assembly_1 and Assembly_2 had 232 Mb (43.0%) and 189 Mb (35.1%) of repeat-masked sequences, respectively (Fig. 2a, Supplementary Table 4).

**Figure 2.**
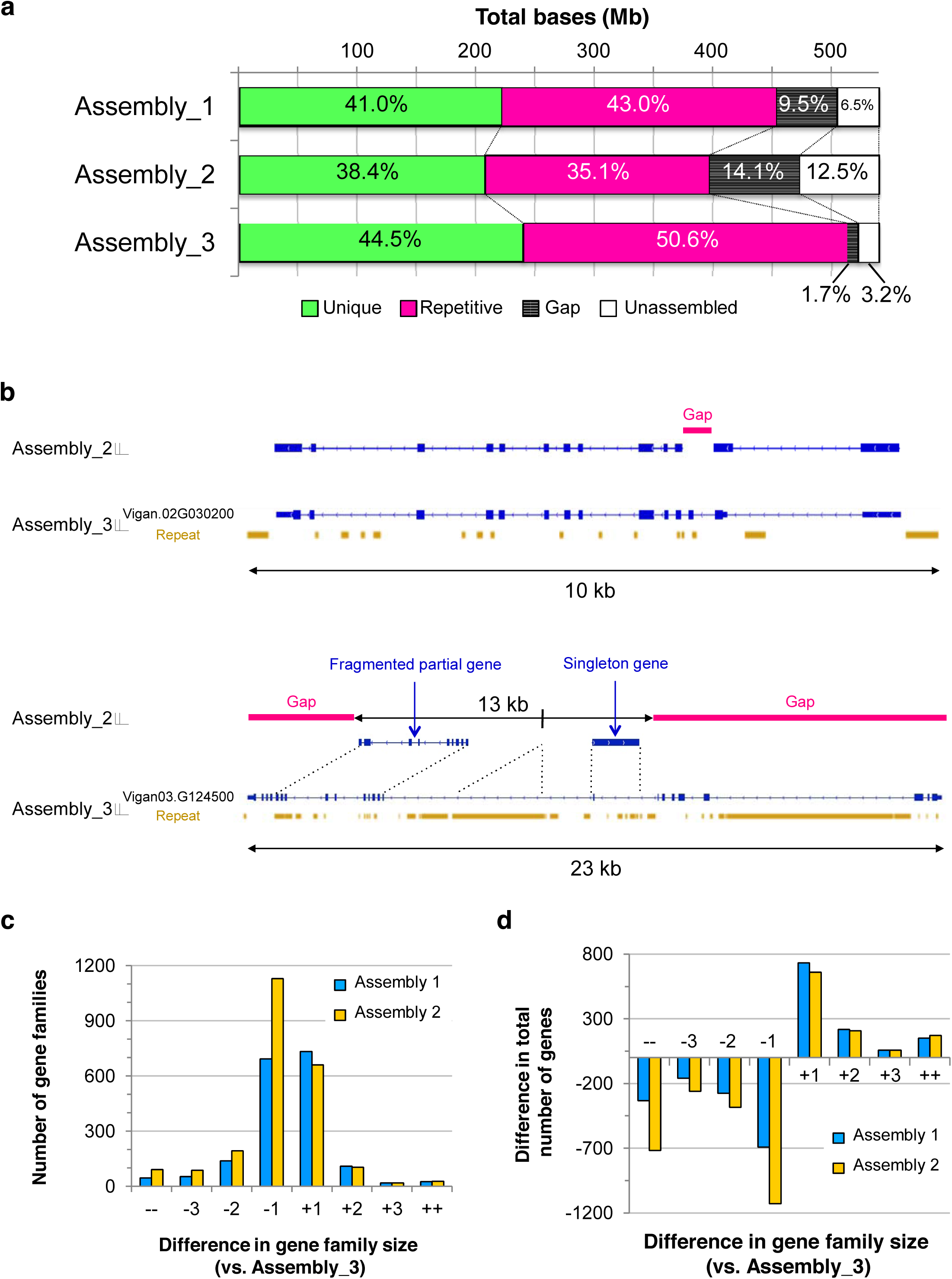
Summary of annotations. (a) The amounts of unique sequences, repetitive sequences, gaps, and unassembled sequences in each assembly. **(b)** Examples of wrong annotations in Assembly_2. At the locus of Vigan.02G030200 (top) in Assembly_3, sequence from the 2nd to the 3rd intron was left as a gap in Assembly_2, leading to fragmentations of this locus. The 23 kb region of the locus Vigan.03G124500 (bottom) was assembled into only a 13 kb contig in Assembly_2, in which both ends of this region were totally unassembled, and a 2 kb region in the 9th intron was missing. In this case, two genes were also annotated, one of which was mostly comprised of intronic sequences. **(c)** Number of gene families with size differences. ++ and -- indicate gene families with differences of more than +4 and □4 in size, respectively. **(d)** Difference in total gene numbers in gene families with size differences.

Interestingly, repeat masking also revealed that the amount of unique (unmasked) sequences greatly varied between assemblies. It was 222 Mb, 200 Mb and 240 Mb in Assembly_1, Assembly_2, and Assembly_3, respectively (Fig. 2a, Supplementary Table 4).

We then performed gene annotation using RNA-seq data of various tissues (Supplementary Table 1) and *ab initio* gene prediction. To compare the quality of the assemblies, we independently annotated all the three assemblies.

As a result, 31,310 protein-coding genes were annotated, of which 30,507 genes (97.5%) were present in the anchored scaffolds in Assembly_3. Assembly_1 and Assembly_2 had 31,153 and 30,187 genes annotated, respectively (Table 1).

To examine the completeness and correctness of the assemblies, we estimated the presence of the core eukaryotic genes (CEGs), a set of 458 genes that are highly conserved in most eukaryotic genomes^24^. The results showed 436, 439 and 447 CEGs in Assembly_1, Assembly_2 and Assembly_3, respectively (Table 1). On anchored scaffolds, 432, 436, and 440 CEGs were present in Assembly_1, Assembly_2, and Assembly_3, respectively (Table 1).

However, the difference in the numbers of annotated genes and CEGs between the assemblies seemed too small, considering the amount of unique sequences in the SGS-based assemblies were 10–20% shorter than Assembly_3. As expected, we found many cases of wrong annotations in the SGS-based assemblies, due to gaps and poor assemblies in coding regions (Fig. 2b).

As described above, incomplete genome assemblies are considered to have collapsed or fragmented genes; thus we clustered all the annotated genes into gene families to compare the number of paralogues between the three assemblies. We took Assembly_3 as a standard and calculated the differences of each family size (Fig. 2c, d).

Of the 15,887 gene families detected, 10,737 families contained the same number of genes in all three assemblies. However, in the gene families with different numbers of genes, Assembly_2 was dominant in the size-reduced families, especially in those with one or more than four genes fewer (Fig. 2c). As a whole, Assembly_2 had 1,499 gene families with reduced size and 811 with increased size, leading to a total of −2,493 and +1,097 genes in the size-reduced and the size-increased families, respectively (Fig. 2d). Assembly_1 showed 887 size-increased gene families (+1,160 genes) and 929 size-reduced families (-1,460 genes) (Fig.2 c, d).

The clustering also revealed that there were more singleton genes (single copy genes that are present only in one assembly) in SGS assemblies. The numbers of singletons were 700, 887, and 557 in Assembly_1, Assembly_2, and Assembly_3, respectively.

### Completeness of the azuki bean genome and other legume genomes

To date, genome sequences of legume crops, including soybean (*Glycine max*)^25^, alfalfa (*Medicago truncatula*)^26^, common bean (*Phaseolus vulgaris*)^15^, chickpea (*Cicer arientinum*)^16^, pigeon pea (*Cajanus cajan*)^17^, mungbean (*V. radiata*)^13^, and azuki bean (*V. angularis*)^14^, have been published (Table 2).

Compared to those genome assemblies, our assembly (Assembly_3) had the best coverage and the least amount of gaps, in both the total assembly and the pseudomolecules (anchored scaffolds) (Table 2).

As shown in Figure 3, the NG graph revealed great variance between the assemblies. The scaffold (pseudomolecule) NG graph showed almost no difference between reference-grade assemblies, including soybean, alfalfa, and common bean, in which Sanger sequencing, BAC libraries, optical mapping, or high-density genetic maps had been integrated. A similar graph was also obtained for Assembly_3, whereas SGS-only draft assemblies, including mungbean, chick pea, pigeon pea, and azuki bean, had a shorter length and coverage of only 20–60% (Fig. 3a).

**Figure 3.**
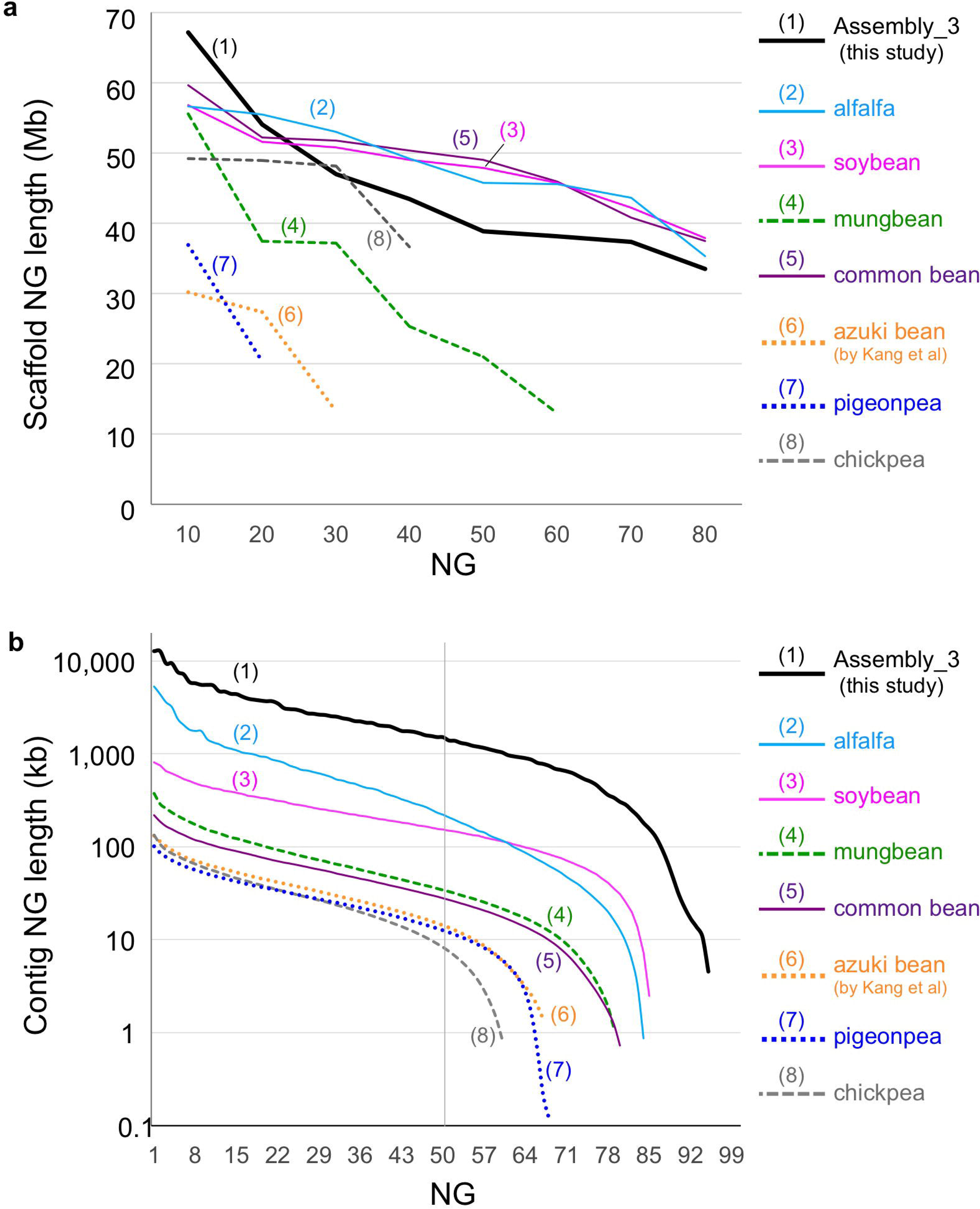
NG graphs of legume genomes of (a) contigs and (b) pseudomolecules. The x-axis indicates NG integers, and the y-axis indicates the calculated NG length in each assembled genome. The vertical line indicates the NG50 contig length. The labels are sorted according to the ranking of contig NG50. The solid lines indicate the reference grade assemblies (total size of anchored scaffolds covering ~80% of genome), whereas broken and dotted lines indicate the draft assemblies (total size of anchored scaffolds covering ~50% and ~30%, respectively).

For contigs, the NG graph revealed that Assembly_3 had the highest contiguity, whereas the SGS-only assemblies were much shorter and were similar to each other (Fig. 3b).

Although all assemblies contained more than 96% of CEGs, the number of CEGs present in the anchored scaffolds ranged from 285 (62.6%) in pigeon pea to 446 (97.4%) in soybean (Table 2). Assembly_3 (440 CEGs) was the second best, following soybean, and was comparable to alfalfa and common bean (439 CEGs) (Table 2).

### Characteristics of the azuki bean genome

Because we obtained a near-complete genome and a high-density genetic map, we could calculate the gene density, amount of repeats, and recombination frequency throughout the whole genome (Fig. 4, Supplementary Fig. 6). Overall, the recombination rate (cM/Mb) positively correlated with gene density, but negatively correlated with repeat density. We also calculated recombination per gene (cM/gene) because the interval lengths between genes are greatly different between gene-rich regions and repeat-rich regions. However, the obtained values were also higher in gene-rich regions than in repeat-rich regions, indicating that recombination is highly suppressed in the putative centromeric and pericentromeric regions (Fig. 4, Supplementary Fig. 6). There were also several regions with no recombinations, other than the centromeric regions, suggesting some structural variation such as inversions between the azuki bean and *V. nepalensis* (Fig. 4, Supplementary Fig. 6).

**Figure 4.**
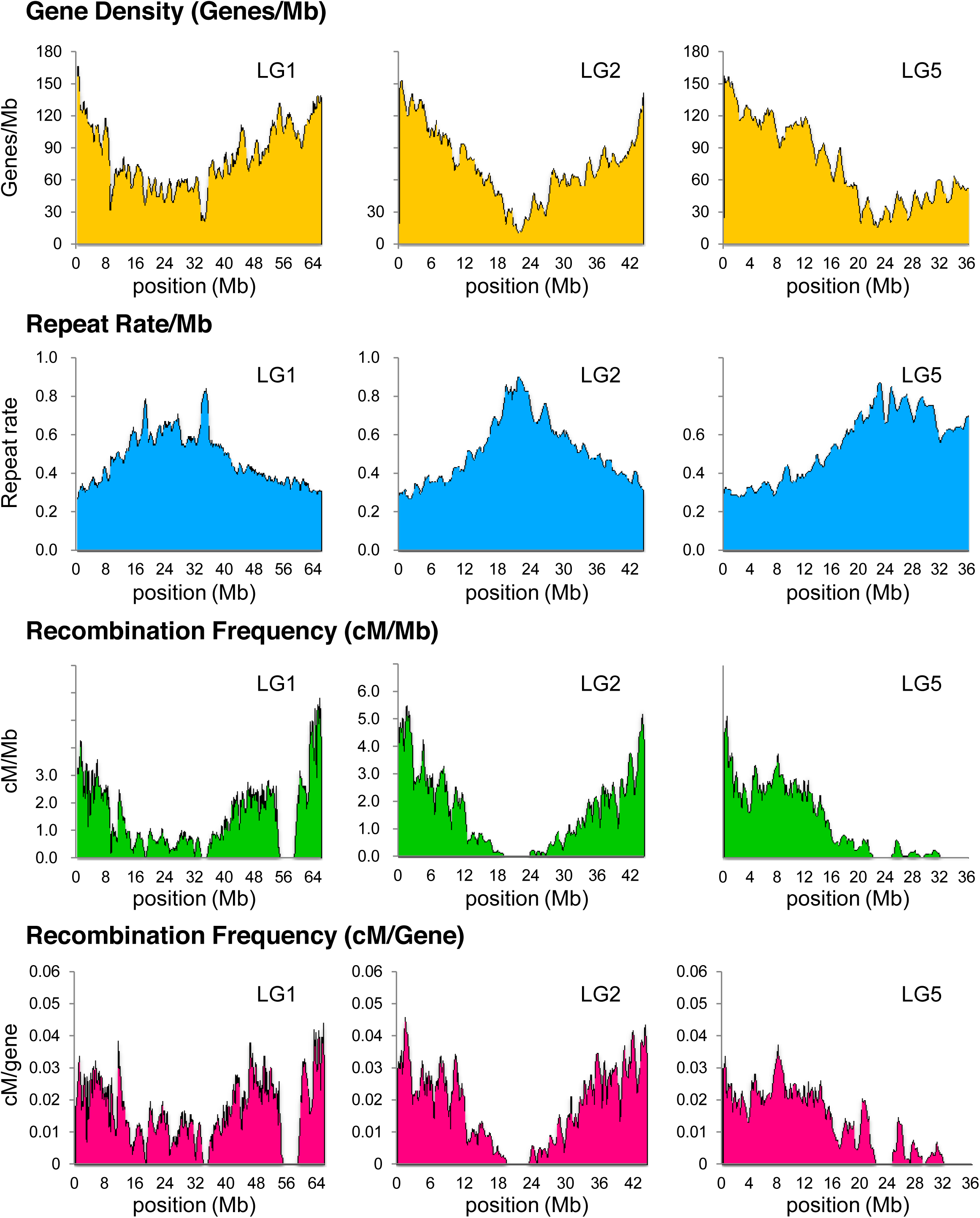
An overview of the azuki bean genome. The x-axis indicates the physical position in Mb in pseudomolecules of LG1, 2, and 5.

## DISCUSSION

Together with SMRT sequencing and a high-density linkage map, we demonstrated that it is possible to obtain a genome assembly of reference-grade, even without fosmid/BAC libraries or optical mapping. The azuki bean genome was successfully assembled into 11 pseudomolecules covering more than 85% of the genome and 97% of the annotated genes, and it was even better, in many aspects, than the existing high-quality legume genomes. In short, compared to most of the sequenced genomes, SMRT sequencing could achieve higher coverage, a smaller amount of gaps, and longer contiguity.

In this study, we *de novo* assembled the azuki bean genome using three major platforms: Roche, Illumina, and PacBio (Table 1, Supplementary Figs. 1, 3, Supplementary Table 1). In Assembly_1, we made extensive efforts to maximize coverage of the azuki bean genome (Supplementary Fig. 1). However, a validation by genetic map revealed it contained lots of assembly or scaffolding errors, which indicated difficulty in optimizing assembly methods. Compared to Roche-based hybrid assembly (Assembly_1), Illumina-only assembly (Assembly_2) was more accurate but lower in coverage. The PacBio (Assembly_3), though it still needed some error correction with short reads, achieved the best contiguity and coverage.

Although the NG graph of scaffolds was not greatly different between Assembly_3 and Assembly_2, the NG graphs of contigs were in a different level (Table 1, Fig. 1). This was also true in a comparison of Assembly_3 with other reference-grade legume genomes, where the NG graphs of scaffolds were similar to each other, but those of contigs were higher in Assembly_3 than in any others (Table 2, Fig. 4). Our NG graph approach also revealed that the pseudomolecules of SGS-only assemblies were much shorter than the estimated genome size of those species (the most was 62% in mungbean) (Table 2, Fig. 4).

To date, many genomes of non-model eukaryotes have been sequenced using Illumina platforms^4^. This is partly because one might expect that even a *de novo* assembly using short reads can capture most of the unique (and thus genic) sequences. Certainly, our results also revealed that large portion of uncaptured sequences in SGS-based assemblies consisted of repeat elements, and the numbers of annotated genes or CEGs were almost the same between the assemblies (Table 1, Fig. 2a, Supplementary Table 4). However, SGS-based assemblies were also missing 20–40 Mb of unique sequence, which accounted for 10–20% of its unique region (Figure 2a). We consider this difference is not too small to be ignored, since SGS-based assemblies contained many gaps and poor assemblies in gene coding sequences, which in turn caused fragmentation, misannotation, and absence of thousands of genes (Fig. 2b-d). Falsely-called singleton genes are also problematic, because they can be easily misunderstood as species-specific genes (Fig. 2b). Thus, the numbers of annotated genes or CEGs are not good indicators for assessing the quality of the assemblies, although they have often been used.

One possible reason for the missing of genic sequences in SGS-based assemblies might be because there are many exons that are not only very short (less than 100 bp) but are neighbored by repeat-rich introns. In such cases, an assembly process only produces contigs that are shorter than minimum threshold, and thus are usually discarded by default.

As such, as argued by Alkan et al^6^ and Denton et al^7^, our results indicate the risk of SGS-only assembly. The existing comparative genomic studies, including those presenting copy number variations (CNVs) and expansion/shrink of some gene families, could reach different conclusions if the reference genomes were again *de novo* assembled using SMRT sequencing technology.

Our results also revealed the importance of having a high-density linkage map when reconstructing pseudomolecules. Although designing SNP arrays is expensive compared to genotype-by-sequencing (GBS) technologies, such as restriction-site associated DNA sequencing (RAD-seq)^27^, it allowed us to design markers at desired intervals and thus to construct a linkage map with unbiased marker distribution (Supplementary Fig. 2). Indeed, we successfully anchored almost 90% of the assembled sequences. Although the GBS approach can easily provide thousands of markers, the linkage maps developed by GBS often have uneven marker distribution^13,14^. As was often observed in linkage maps using amplified-restriction fragment length polymorphism (AFLP) markers^28,29^, there are clusters of many RAD-seq markers and large gaps of more than 10 cM. In the draft azuki bean genome^14^, the linkage map was constructed by RAD-seq, and the anchored scaffolds covered only one-third of the genome. In contrast, when we anchored their scaffolds using our linkage map, the total length of the anchored scaffolds almost doubled.

Thus, GBS is probably not the best approach for reconstructing pseudomolecules of *de novo* assembled sequences, unless one is resequencing the whole genomes of the mapping population. It might also be important to choose appropriate parents, with whom little segregation distortion is observed in later generations.

Although we emphasized the strength of SMRT sequencing, it is not perfect. In bacterial genome assemblies, relatively higher error rates can be overcome and a phred score of Q60 is achieved with ~50× coverage of SMRT sequencing data^30^. However, in the azuki bean genome, 50× coverage could not achieve such high accuracy. The nucleotides with low phred scores were greatly enriched at homopolymer sites, which are present much less in bacterial genomes. All the low-score nucleotides conflicted with Illumina reads turned out to be wrong, as demonstrated by Sanger sequencing (Supplementary Table 2).

As such, assembling eukaryotic genomes with a SMRT sequencer would still need error correction by short reads. Of note, if not corrected, the indel errors in Assembly_3 would break the coding frames of more than 1,000 annotated genes.

Because we performed a linkage analysis with 1,000 F2 plants, the genetic map had a resolution of 0.1 cM. This high-resolution genetic map enabled us to estimate the recombination frequency per gene (cM/gene) throughout the whole genome (Fig. 4, Supplementary Fig. 6). With this value, one can directly predict the population size that is necessary to isolate target genes in a desired region. If a target gene is present in a region of 0.05 cM/gene, it means a mapping population of 2,000 plants will be enough. If it is in a 0.01 cM/gene region, a population of 10,000 plants will be required.

Plant geneticists have empirically noticed that genes in euchromatic regions are much easier to clone than those in heterochromatic, repeat-rich regions, despite the physical intervals between genes being much longer. For example, the soybean *Pdh1* gene is present in a euchromatic region and was isolated with a mapping population of 2,535 plants^31^, whereas the *E1* gene in a pericentromeric region required approximately 14,000 plants to obtain a recombinant between the neighboring genes^32^. As such, a clear correlation between gene density and recombination rate per gene in our results strongly supports such notions. We hope it will greatly support breeding activities including gene cloning and MAS in azuki bean.

## METHODS

Methods and any associated references are available in the online version of the paper.

**Accession Codes.** Raw sequence data generated in this study are available at DDBJ under the BioProject ID PRJDB3778. The data from *V. nepalensis* is available under DDBJ BioProject ID PRJDB3779. Genome assembly and annotation are available at http://viggs.dna.affrc.go.jp/download.

## ACKNOWLEDGEMENT

We are grateful for Dr. Kazuaki Hashimoto, Dr. Kenichi Dedachi and Mr. Ken Osaki in TOMMY DIGITAL BIOLOGY CO., LTD and Dr. Stephen Turner in Pacific Biosciences of California, Inc. for their kind and useful advice on technical issues. We also appreciate Dr. Yuki Monden at Okayama University and Dr. Shinpei Kawaoka at Advanced Telecommunications Research Institute International for valuable academic discussion. This research project was funded by the Scientific Technique Research Promotion Program for Agriculture, Forestry, Fisheries, and Food industry.

## AUTHOR CONTRIBUTIONS

HS did the whole genome assemblies and comparative analysis, and wrote the manuscript. KN conceived the study, participated in its design, analyses, and coordination, and wrote the manuscript. EOT, YT, KI, and CM cultivated F2 plants, and performed genotyping and linkage map construction. KS performed sequencing and wrote the manuscript. KT, AS, and MS performed sequencing. TH managed sequencing and participated in study design and coordination. TI performed the comparative analysis between assemblies and helped draft the manuscript. AK developed and cultivated F2 plants, and performed linkage analysis. NT participated in study design and coordination, and helped draft the manuscript. All authors read and approved the final manuscript.

## COMPETING FINANCIAL INTERESTS

The authors declare that they have no competing interests.

## ONLINE METHODS

**Plant materials and DNA/RNA extraction.** Seeds from *V. angularis* cv. ‘Shumari’ were provided by the Tokachi Agricultural Experiment Station of Hokkaido Research Organization, Memuro, Hokkaido, Japan, and seeds from *V. nepalensis* (JP107881) were provided by the NIAS Genebank, Japan. The seeds were used for genome sequencing and transcriptome analysis. We grew plants in a greenhouse in Tsukuba, Japan and extracted DNA from unexpanded leaves using the CTAB method [38], followed by further purification using the Genome Tip 100/G (Qiagen K.K., Tokyo, Japan).

We extracted RNA using the RNeasy Plant Mini Kit (Qiagen) from shoots, leaves, stems, roots, and root nodules of 2-week-old plants and the flowers of 2-month-old plants. After harvesting seeds, we removed the seed coat from dried seeds, separated axes and cotyledons, and extracted RNA using the phenol/SDS method^33^.

For genetic linkage map construction, we crossed *V. nepalensis* (JP107881 in NIAS Genebank, Japan) to *V. angularis* cv. ‘Erimoshouzu’ (JP37752) and obtained 1,000 F2 seeds. We grew the F2 plants in an incubator, MIR-253 (Sanyo), for a week and extracted DNA using the CTAB method^34^.

**DNA sequencing.** For the Roche GS Titanium and FLX+ platform, construction of single-end and paired-end libraries and sequencing were all provided as a custom service of Beckman Coulter Genomics (Danvers, MA, USA). In total, we performed 19 runs of the single-end library using the GS FLX+ and four runs of the 3 kb mate-pair library, four runs of the 8 kb mate-pair library, and three runs of the 20 kb mate-pair library using the GS Titanium.

For the Illumina HiSeq 2000 platform, library construction and sequencing was provided as a custom service of Eurofins MWG GmbH (Ebersberg, Germany). Sequencing libraries included a paired-end library of 300 bp inserts and mate-pair libraries of 3 kb, 8 kb, 20 kb, and 40 kb inserts. One lane of the flow cell was used for each sequencing library.

For the Illumina HiSeq 2500 platform, library construction and sequencing was provided as a custom service of Macrogen Inc (Seoul, South Korea). A paired-end library of 270 bp inserts was constructed, and one lane of the flow cell was used for sequencing.

After sequencing, adapter and low-quality sequences were trimmed off using Trimmomatic ver. 0.32^35^.

For the PacBio RS II platform, the extracted DNA was sheared into 20 kb fragments using g-TUBE (Covaris, MA, USA) and converted into 20 kb SMRTbell template libraries. The library was size selected for a lower cutoff of 7 kb using BluePippin (Sage Science, MA, USA). Sequencing was performed on the PacBio RS II using P5 polymerase binding and C3 sequencing kits with 180 min acquisition. In total, 79 SMRT cells were sequenced, and about 27.6 Gb of reads were produced.

### *De novo* assembly of the *V. angularis* genome

**Assembly_1.** Roche 454 reads were assembled using the Celera Assembler ver. 7.036 with the following options: utgErrorRate = 0.015, ovlErrorRate = 0.03, cnsErrorRate = 0.03, and cgwErrorRate = 0.05. To correct errors in the contigs, we first mapped Illumina paired-end reads to the contigs using the Burrows-Wheeler Aligner (BWA) ver. 0.6.2^37^ under the default setting. We further refined the alignments around indels using the IndelRealigner of the Genome Analysis Toolkit (GATK) ver. 1.3-25^38^ and discarded the putative polymerase chain reaction (PCR) duplicates using the MarkDuplicates of Picard ver. 1.63 (http://picard.sourceforge.net/). Then, substitutions and indels were detected using the mpileup of Samtools ver. 0.1.18^39^. Only variant sites with ≥10 reads that have ≥30 phred score, ≥70% frequency, and no strand bias (*p* > 0.01) were selected as errors. Finally, we replaced the error sites with Illumina sequences. The error correction step was repeated three times. Low-quality sequences at the edges of the error-corrected contigs (ecContigs) were trimmed off using the trimFastqByQVWindow.py in SMRT Analysis (http://www.pacb.com/devnet/) ver. 1.4.0 with the —qvCut = 54.5 and –minSeqLen = 200 options.

To further extend the ecContigs, we mapped the Illumina reads to the ecContigs and assembled unmapped Illumina reads, combining them with the ecContigs using the String Graph Assembler (SGA) ver. 0.9.35^40^. In assembling using SGA, first, duplicated sequences and reads with low-frequency k-mers in the Illumina reads were filtered using the default setting. Because a large number of the ecContigs were expected to contain unique k-mers, Illumina reads and ecContigs were merged and filtered using the “sga filter --no-kmer-check” option to avoid discarding ecContigs containing unique k-mers. Final assembling was done with the “sga assemble -m 75 -d 0.4 -g 0.1 -r 30 -l 200” command. To screen organelle-originating contigs, we conducted BLASTN searches against the *V. angularis* organelle sequences^41^ and discarded the ecContigs that had ≥98% identity and less than 200 bp of unmatched sequences. We further discarded the contigs matching to any non-plant genomic sequences with ≥90% identity and ≥95% sequence coverage in the nt database of NCBI. Scaffolds were constructed using SSPACE ver. 2.0^42^ with the -z 200 and -k 5 options using both Roche paired-end and Illumina mate-pair reads.

To close the sequence gaps in the scaffolds, we first corrected sequencing errors in PacBio (P4-C2) reads using Roche 454 reads using the pacBioToCA command implemented in SMRT Analysis with these options: utgErrorRate = 0.25, utgErrorLimit = 4.5, cnsErrorRate = 0.25, cgwErrorRate = 0.25, and ovlErrorRate = 0.25. Then we ran PBJelly ver. 12.9.14^43^ using the error-corrected PacBio reads with the following options: blasr -minMatch 8, -minPctIdentity 70, -bestn 8, -nCandidates 30, -maxScore -500, -nproc 12, and -noSplitSubreads.

After closing the sequence gaps, we split the scaffolds into contigs, conducted SSPACE again, and obtained the final scaffolds.

**Assembly_2.** Illumina reads were assembled using the ALLPATHS-LG (47212)^23^ program under the default setting. Organelle sequences were discarded in the same manner as in the construction of Assembly_1. Any scaffolds having BLASTN hits to non-plant organisms with ≥80% identity across ≥200 bp were also discarded.

**Assembly_3.** PacBio (P5-C3) reads were corrected using Sprai ver. 0.9.5.1.3^44^, and the longest 25× error-corrected reads were run with Celera Assembler ver. 8.2beta under the utgErrorRate = 0.02, utgErrorLimit = 4.5, cnsErrorRate = 0.25, cgwErrorRate = 0.25, and ovlErrorRate = 0.02 options. Contigs that were aligned to other contigs with over 98% of the sequence and with over 98% identity were excluded. Assembled contigs were polished by Quiver in SMRT Analysis ver. 2.2.0. To correct indel errors in the contigs, first, we mapped the Illumina paired-end reads to the contigs using BWA-MEM ver. 0.7.9. After conducting local realignment and discarding PCR duplicates as described above, we detected indels using the HaplotypeCaller of GATK ver. 3.2^38^ and selected only reliable indels using VariantFiltration under the following settings: --filterExpression DP < 10 || DP > 100 || QD < 2.0 || FS > 60.0 || MQ < 40.0. Then, we selected only homozygous indels and replaced them with Illumina sequences using the FastaAlternateReferenceMaker of GATK ver. 3.2^38^.

To assess the accuracy of the assembly, we mapped the probe sequences of the 6,000 SNP markers (see the ‘*Linkage analysis*’ section below), as well as Illumina reads, to the contigs. If SNP markers in one or more LGs were mapped to a single contig, we detected and discarded misassembled regions by manually inspecting the read alignments. Organelle and possibly contaminated contigs were discarded as described above.

The assembled contigs were scaffolded using SGA ver. 0.10.13^40^ on Illumina mate-pair reads. If sequence gaps with negative sizes were detected, we aligned 10 kb of flanking sequences and connected the flanking contigs if ≥500 bp were aligned with ≥95% identity. We ran PBJelly2 three times to close as many sequence gaps as possible using the error-corrected PacBio (P5-C3) reads. Finally, we corrected indel-type errors as described above and obtained the final scaffolds.

**Linkage analysis.** We constructed a paired-end library of 300 bp inserts and resequenced *V. nepalensis* using the HiSeq 2000 (Illumina). Library construction and sequencing was provided as a custom service of Beckman Coulter Genomics. To construct the initial set of SNPs, we first mapped the reads onto Assembly_1 using BWA^37^ and extracted single nucleotide polymorphisms in the same manner as in constructing Assembly_1. We selected homozygous SNPs with ≥10 covering reads that have ≥30 phred score, ≥90% frequency, and no strand bias (*p* > 0.01). Then, we further selected SNPs where the flanking 60 bp of sequence contained no other SNPs and had no BLASTN hits on Assembly_1 (E value < 1.0e^-10^). To make sure that the flanking sequences are fixed and are not heterozygous in *V. angularis*, we mapped the Illumina paired-end reads from *V. angularis* to Assembly_1 and selected only SNPs where every site of the flanking 60 bp of sequence was called as the same homozygous genotype as Assembly_1.

Of these, we selected 6,000 SNPs from the 1,036 longest scaffolds of Assembly_1 and designed probes for the infinium assay (Illumina)^45^ such that the selected SNPs were as evenly distributed as possible.

Genotyping using the infinium assay was provided as a custom service of Medical & Biological Laboratories Co. Ltd (Nagoya, Japan). We validated the genotype data by R/qtl^46^, removed samples and markers with more than 10% missing data, and grouped the markers into 11 LGs. Marker orders and distances were estimated by Antmap^47^.

**Reconstructing the pseudomolecules.** Probe sequences for the 6,000 SNP markers were mapped to the Assembly_3 scaffold sequences, and only uniquely and identically mapped SNPs were used to order the scaffolds along the linkage map. If a scaffold was anchored by only one SNP marker, or all markers on a scaffold were tightly linked to each other, it was impossible to determine the orientation of the scaffold. Therefore, we tentatively determined the orientation of such scaffolds from the synteny information between Assembly_3 and *P. vulgaris* generated by MUMmer ver. 3.23^48^. The size of each gap was estimated from the intervals between the nearest two SNP markers on each of the adjacent sequences. If it was impossible to estimate the size, 1,000 Ns were inserted. Finally, we assigned the chromosome numbers in accordance with the previous study^49^ by mapping the probe sequences of the SSR markers.

**Repeat prediction.** A repeat sequence library was first constructed *de novo* by RepeatModeler ver. 1.0.8 [http://www.repeatmasker.org], and then combined with the MIPS Repeat Element Database ver. 9.3^50^. We detected the repetitive elements using CENSOR^51^.

### Validation of the nucleotide-level accuracy of PacBio assembly

We downloaded the raw data of *A. thaliana* Ler-0 from the PacBio DevNet (https://github.com/PacificBiosciences/DevNet/wiki/) and conducted *de novo* assembly by the same manner as described above. Besides, we obtained the genome assembly of *A. thaliana* Ler-1 from 1001 Genomes Data Center (http://1001genomes.org/datacenter/). The Ler-1 assembly was derived by assembling the Illumina sequences. Then we constructed the pairwise genome alignment of the two assemblies and detected discrepant sites by MUMmer. For *D. melanogaster* assemblies, we obtained PacBio assembly from PacBio Devnet and Sanger-based reference assembly from Ensembl^52^ and detected discrepancies.

**RNA-Seq analysis** Library construction and sequencing was provided as a custom service of Beckman Coulter Genomics. Two lanes of the Illumina HiSeq 2000 were used to sequence eight libraries.

### Gene prediction

**TopHat and Cufflinks.** Gene structures were predicted using TopHat ver. 2.0.13^53^ and Cufflinks ver. 2.2.1^54^ with the --min-intron-length 50 and --max-intron-length 10000 options for TopHat. Open reading frames (ORFs) were predicted using TransDecoder ver. r20140704 and Trinotate ver. r20140708^55^. The ORF with the highest identity and amino acid length was selected as the best ORF for each transcript. Transcripts where no ORF was predicted were annotated as non-coding genes.

**PASA2.** Besides TopHat and Cufflinks, we predicted gene structures using genome-guided and *de novo* RNA-Seq assembly approaches using Trinity ver. r20140717^54^ and the PASA pipeline^56,57^. Before assembling the RNA-Seq reads *de novo*, we merged the eight libraries and conducted *in silico* normalization implemented in Trinity to reduce the data size. Genome-guided and *de novo* assembly was done using Trinity with the --genome_guided_max_intron 10000 option and the default setting for genome-guided and *de novo* assembly, respectively. The PASA pipeline was run under the default setting. As described above, an ORF was predicted for each transcript, whereas transcripts with no predicted ORF were annotated as non-coding genes.

**MEGANTE.** We carried out gene prediction using MEGANTE^58^ with the parameters of *V. unguiculata*. Genes with greater than 40% of their transcribed and exonic regions masked as repeat sequences were discarded.

**Merging the gene structures.** First, gene structures predicted by TopHat and Cufflinks, and the PASA pipeline were clustered on the genome assembly, and the representative structure with the longest ORF was selected for each locus. Then, MEGANTE-predicted genes in unannotated regions were added. To screen transposon-related genes, we conducted BLASTP searches against UniProt release 2014_07 and RefSeq release 66, and discarded genes matching any transposon-related proteins with an E value <1.0e^-10^. Additionally, genes containing transposon-related functional domains were removed using InterProScan ver. 5.7-48.0^59^. The remaining genes were subjected to expression validation analysis with the RNA-Seq data. RNA-Seq reads from the eight libraries were mapped separately using TopHat. The expression level was estimated by Cuffquant for each locus and then normalized by Cuffnorm^60^. A gene was discarded if its expression level was zero FPKM in all eight libraries and it did not have any homologs in the BLASTP result. The remaining genes were selected as the final gene set.

We carried out the whole annotation procedure described above for Assembly_1, 2, and 3.

**Gene family prediction.** Protein sequences predicted on all three assemblies were merged and subjected to all-to-all BLASTP searches with the -evalue 1.0e-10 option. Gene families were predicted using mclblastline^61^ with the --blast-score = b, --blast-sort=a, --blast-m9, --blast-bcut=5, and --mcl-I=2.5 options.

